# The Prevalence of Litter Foraging Among UK Birds Lessons Learned From A Citizen Science Project

**DOI:** 10.1101/2021.11.16.468840

**Authors:** Sabrina Schalz, Warren D. Horrod-Wilson, Keir Chauhan

## Abstract

Anthropogenic food sources, such as litter, are readily available to birds in urban areas. As an overly anthropogenic diet can have negative health consequences for wildlife, monitoring the frequency of foraging in litter compared to natural food resources can be an important component of wildlife conservation efforts. To understand how common litter foraging is among different bird species, we conducted a citizen science project with volunteers across the UK. Through this project, we also tried to engage people in birdwatching and bird surveys who would not normally participate in these activities. A particular focus was on recruiting respondents from underrepresented groups, and the methodology was designed to accommodate people without any background knowledge of birds. While we did not receive enough observations to draw conclusions about the litter foraging rates of different bird species across the UK, we report the submissions we received, as well as the demographic composition of the volunteer group. We successfully reached volunteers who did not participate in birdwatching or BTO bird surveys before, as well as some young volunteers, but were unsuccessful in reaching respondents from diverse socioeconomic backgrounds. We hope that the successes and failures of our methods reported here can be useful to others designing citizen science studies, so that birdwatching and volunteering for bird surveys will be equally accessible to all in the future.

## 2. Introduction

Urbanisation and human presence have provided novel foraging opportunities for many bird species (Isaksson, 2018): Carrion Crows (*Corvus corone*) have been found to aggregate in areas rich in anthropogenic food sources, such as waste disposal sites (Preininger et al., 2019). House Sparrows (*Passer domesticus*) are more abundant in areas where there is access to rubbish containers, compared to areas where rubbish is stored underground or managed more effectively (Bernat-Ponce et al., 2021). Feral pigeon (*Columba livia*) populations have become dependent on anthropogenic food sources, with a sharp decline in population sizes occurring during the COVID-19 circuit-breaker lockdown period in Singapore (Soh et al., 2021). Other than waste disposal sites, birds generally forage in areas where human activity occurs, such as picnic areas (Kurosawa et al., 2003), where they may directly take food from humans (Kurosawa 1999, as cited in Kurosawa et al., 2003).

But easy access to anthropogenic food comes at a cost: Innutritious diet has been linked to leucism in urban blackbirds (*Turdus merula*) (Izquierdo et al., 2018). In urban Red-Winged Starlings (*Onychognathus morio*) nestlings fed a diet comprised of a higher proportion of anthropogenic food grew at a significantly slower rate compared to that of nestlings fed a higher proportion of naturally occurring food (Catto et al., 2021). Adult birds who consumed a higher proportion of anthropogenic food had a significantly greater body mass compared to that of individuals which consumed a higher proportion of naturally occurring foods (Catto et al., 2021). A study conducted on urban American Crow (*Corvus brachyrhynchos*) nestlings found that they had higher levels of cholesterol compared to their rural counterparts, though evidence as to whether cholesterol negatively or positively impacts avian life is mixed (Townsend et al., 2019). Birds have also been observed to eat cigarette butts which contain toxic chemicals and plastics, which may consequently impact health and survival (Torkashvand et al., 2020). Plastic has been found in the faeces of urban black vultures (*Coragyps atratus*) and turkey vultures (*Cathartes aura*), but almost none was found in that of Andean condors (*Vultur gryphus*), which avoid urban habitats (Balleko et al., 2021).

Organisations such as the British Trust of Ornithology (BTO) are heavily reliant on the work of citizen scientists to get a national picture of our bird populations. Without continued involvement in their citizen science they would be unable to provide detailed analysis of changes to the UK’s bird population. In doing so, the BTO has helped to establish a community around birding; whether that’s working with bird observatories, supporting young people, or getting people outside and using their green spaces. Citizen science’s impacts on mental health and wellbeing for many people and particularly those of young people is of paramount importance (White et al., 2019). However, more work still needs to be done to encourage a more diverse population of birders participating in bird surveys and to ensure their safety when joining. Women, young people, people of colour, people without a background in higher education, and people in lower income households remain minorities within birding (US Fish & Wildlife Service, 2009).

The first aim of this study was to examine the frequency of foraging in anthropogenic resources such as litter and food waste across the UK, and across different species. Understanding the prevalence of litter foraging along the urban-rural gradient, as well as potential differences between species, is an important component of wildlife conservation work. The second aim was to develop a survey that is accessible to volunteers from the general public who don’t have any previous knowledge about birds. Projects such as this can help engage more people with the natural world, and in particular the birds in front of their doorsteps. While not necessarily collecting much usable data and not a solution to other deterring factors such as racism, sexism, and financial constraints, this project seeks to help engage underrepresented groups by reducing the barrier of required prior knowledge and prior established contact with the bird community through our design of the questionnaire and use of recruitment channels. Recruitment of a citizen science sample not only provides the opportunity to engage different communities in bird watching, but it also presents the opportunity to educate people and encourage behavioural change initiatives (Van Noordwijk et al., 2021), such as reduced rates of littering. We focused on reaching groups currently underrepresented in birdwatching, in particular with regards to age and socioeconomic backgrounds.

## 3. Materials and Method

### 3.1 Subjects and Recruitment

Participants (N = 55) were UK residents (see demographic summary under 4.2). A total of 254 clicks on the survey link were recorded.

To reach a diverse and large group of potential participants, we used a variety of communication channels for recruitment. Emails sent to approximately 1,000 community groups across the UK interested in conservation was the most successful method. Twitter (17,9012 views and 51 link clicks on the main tweet during two months of data collection), posts in public Facebook groups, BTO newsletters, emails to approximately 200 London youth groups, and approximately 50 flyers with QR codes put up in London were moderately successful. The office of the Mayor of London has also helped to distribute the survey among their contacts. A TikTok video (preferred platform for under 25s, 17 likes and 536 views during the second month of data collection), emails to summer camps, schools and 250 student unions across the UK were not successful.

### 3.2 Questionnaire

The survey was presented online in Qualtrics from 26^th^ June to 26^th^ August 2021 (see questionnaire in appendix A). A common barrier in citizen science bird surveys is that most require bird ID skills (personal observation). We designed the questionnaire in a way that it requires no previous knowledge of birds so that more people without a background in birding could participate. Demographic questions were included to evaluate the success levels of reaching respondents from different backgrounds. Respondents were also asked to make up a username for themselves so that we could track multiple participations. The survey was approved by the Middlesex Psychology Ethics Committee and all participants gave informed consent before beginning the survey.

### 3.3 Analysis

61 responses were included in the analysis. Two observations had been excluded as they had taken place outside the UK. Blue tit, great tit, robin, goldfinch, greenfinch, sparrow, and thrush were pooled into the umbrella term “Garden bird”. Mallard, goose, jackdaw, rook, siskin, treecreeper, and green woodpecker were excluded from the group analysis as there was only one observation of each, but their data is reported in appendix B (see table 1). The other species were grouped into blackbirds, starlings, magpies, pigeons, crows, and gulls (each with seven to 13 observations). The share of human and natural resources, as well as the share of foraging habitats are reported for each group. Demographic distributions of participants are also reported to examine the outreach success of this project.

## 4. Results

### 4.1 Foraging

Observations were made in 37 different cities and towns: 47 observations in England, five in Scotland, and five in Wales in total (see appendix B table 2 for list of cities and towns). 26 observations were made in the afternoon (PM), 33 in the morning (AM). 13 foraging events lasted less than one minute, 14 lasted approximately one minute, 24 approximately five minutes and seven lasted 10 minutes or longer. 11 observations were of single birds foraging alone, 47 were in a group of conspecifics or heterospecifics.

29 birds were spotted handling human food, 32 handling nature food. Human foods included bread, fish heads and leftovers, while natural foods included a discarded apple, commercial bird food and nuts. While these items were provided by humans, they are indiscernible from naturally found apples, seeds and nuts and were therefore counted as natural. Bread and fish leftovers do not naturally occur in the corresponding habitats (though there is obviously some overlap between human and natural foods, since humans also consume the latter and don’t rely exclusively on processed foods).

There were considerable differences between species groups: garden birds, blackbirds and starlings were rarely reported to take human food, while crows and gulls mostly took human food, and magpies and pigeons took a combination of natural and human foods (see figure 1).

**Figure 1:**
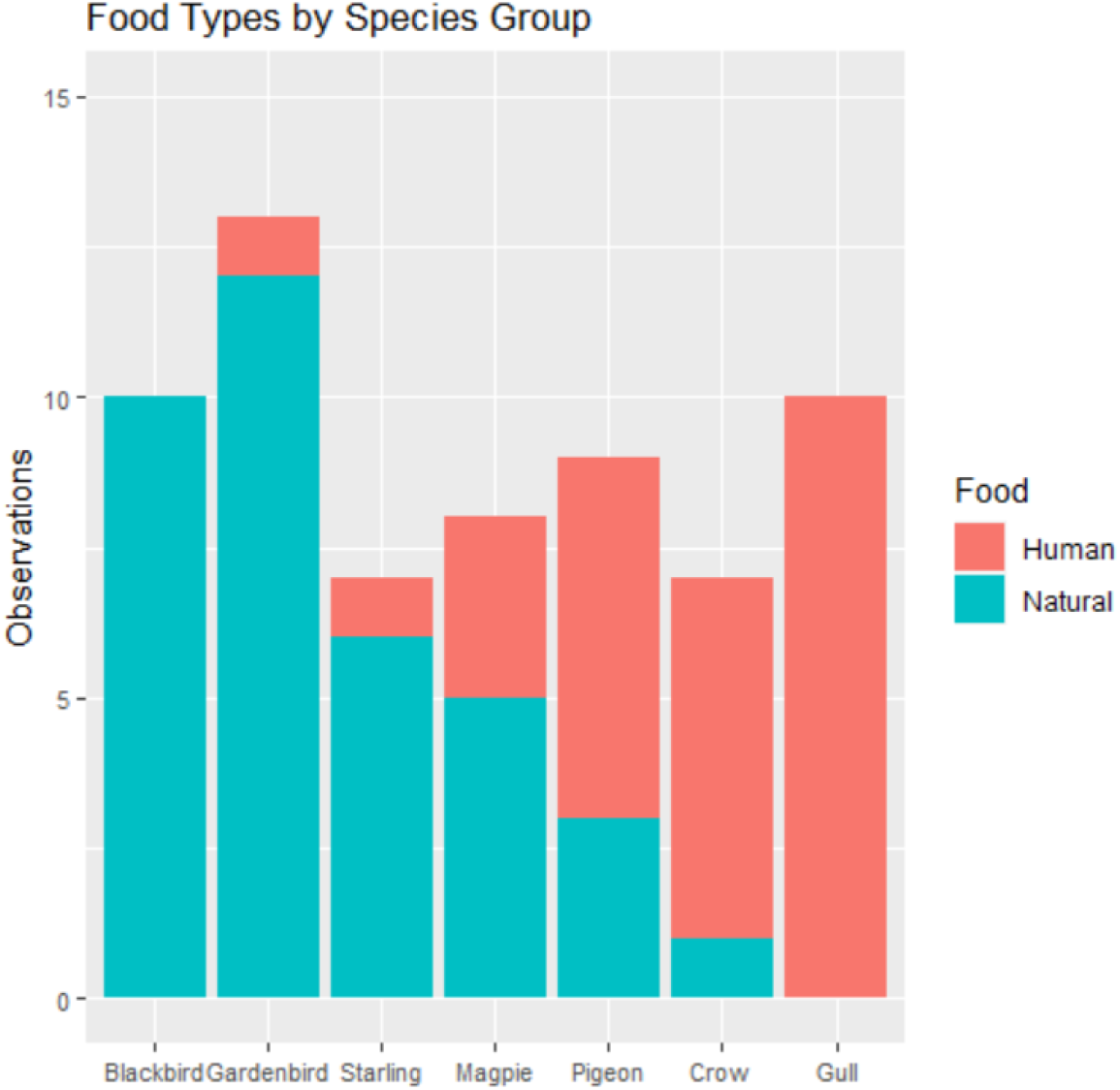
Food types (human or natural) by species group.

Those species preferring human food sources had more diverse foraging locations while those relying exclusively or almost exclusively on natural foods were almost entirely seen in gardens (see figure 2).

**Figure 2:**
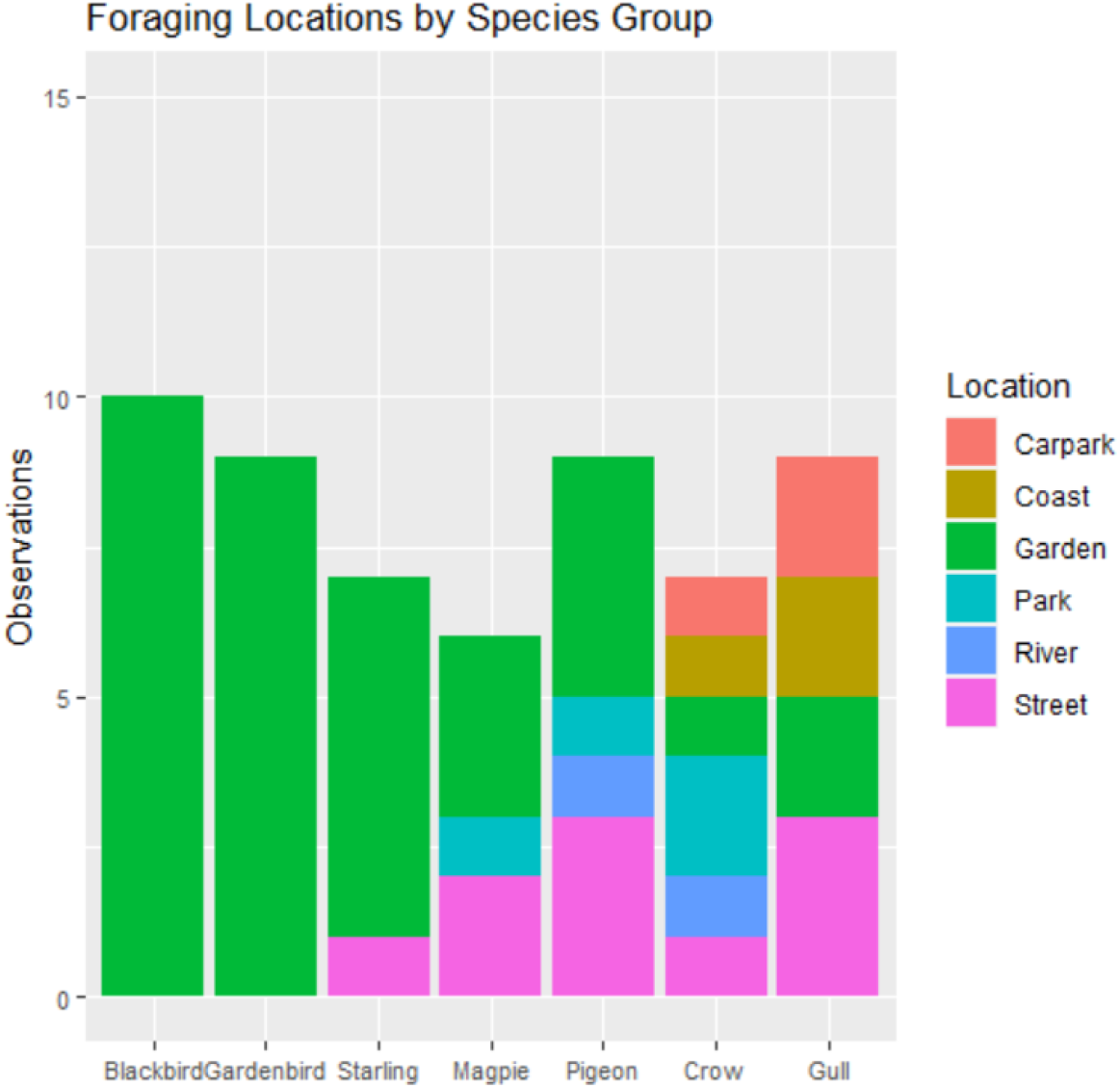
Foraging locations/habitat types by species group, only including observations were location was specified.

Across species, the foraging locations between human foods and natural foods only differed with regards to human food coming from bins and natural foods coming from bird feeders instead. Both food types were equally distributed with regards to the other locations (provided by a person, found on the ground). Only crows, magpies, pigeons and gulls were reported to forage in bins.

### 4.2 Participants

29 participants self-reported their gender as female, 22 as male, one as non-binary and three did not specify. The mean age of respondents was 51.7 (SD = 19.5, 2 participants did not report their age, see figure 3).

**Figure 3:**
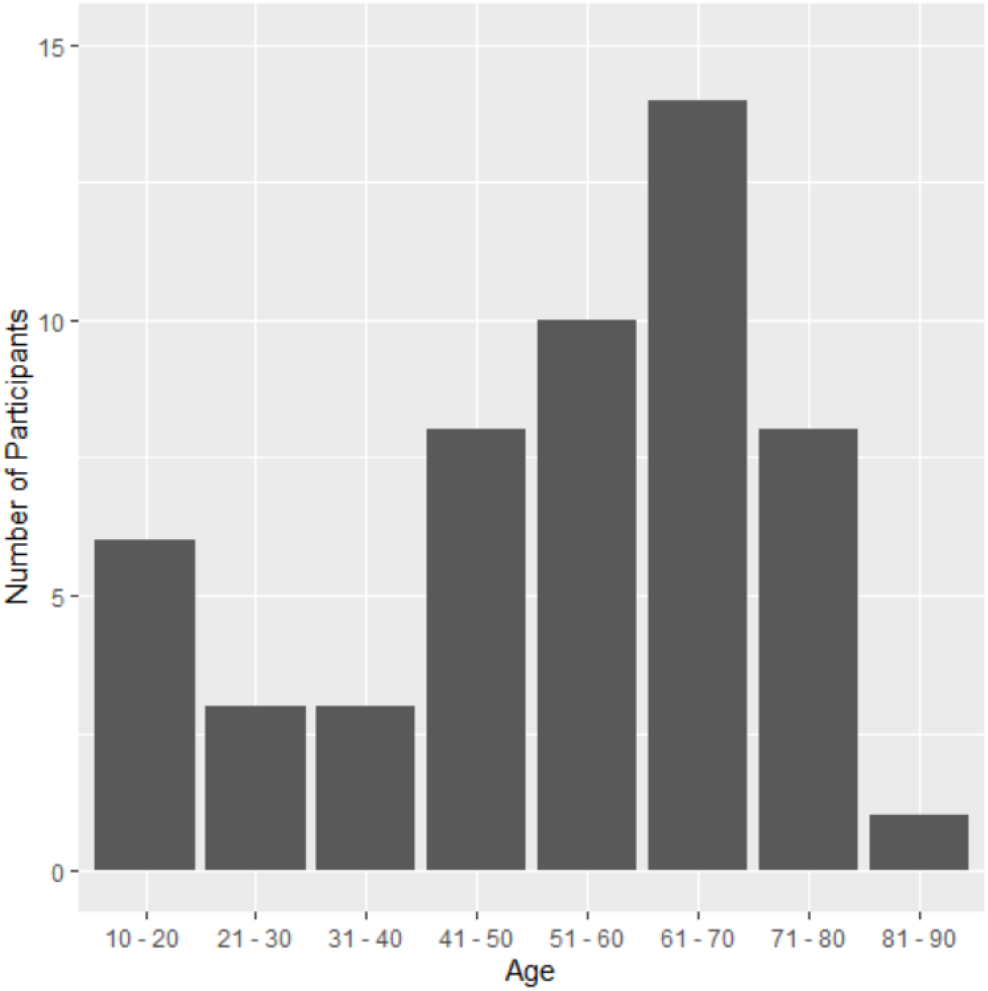
Age distribution of participants.

Four respondents participated multiple times. One respondent changed their answer from “never birdwatching” to “birdwatching less than once a month” in their second participation. 13 respondents made use of the option to use common names for birds instead of their species name (such as “seagull”). Another person described the appearance of a bird instead of naming it.

45 respondents self-identified as British, one as Caribbean, one as British-Caribbean, one as Black British, one as Dutch, one as European, and one as Polish. Most volunteers had completed either college or a university degree and have a yearly household income of £20,000 to £40,000 (see figure 4).

**Figure 4:**
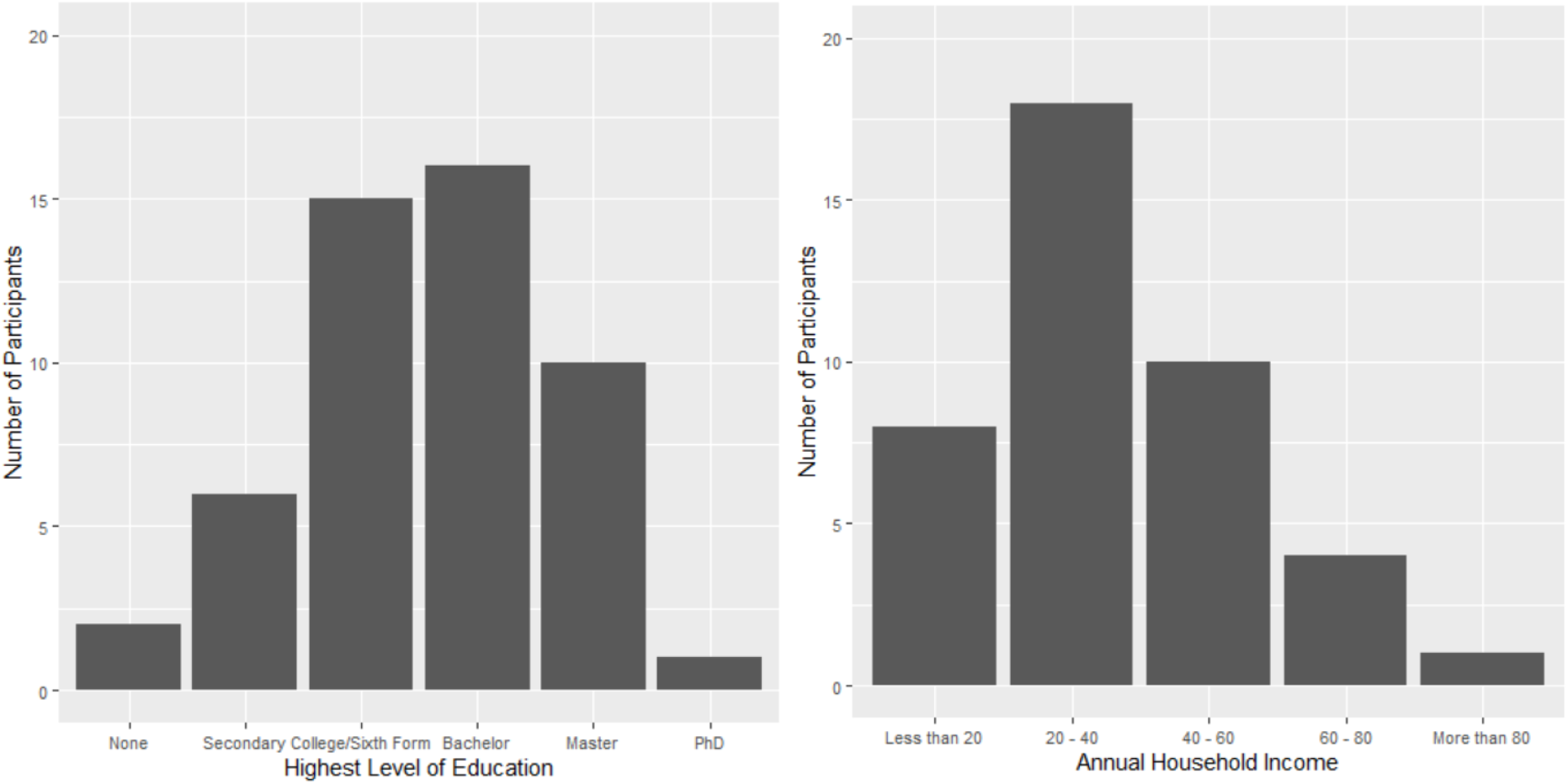
Distribution of highest level of education achieved by participants (left) and annual household income of participants in thousand pounds units (right).

The majority of volunteers notice birds all the time (N = 39) or most of the time (N = 12), and report liking birds a lot (N = 41). However, only half of volunteers engage in birdwatching regularly and a considerable share never go birdwatching (see figure 5). Four respondents had previously volunteered for the BTO, 49 had not.

**Figure 5:**
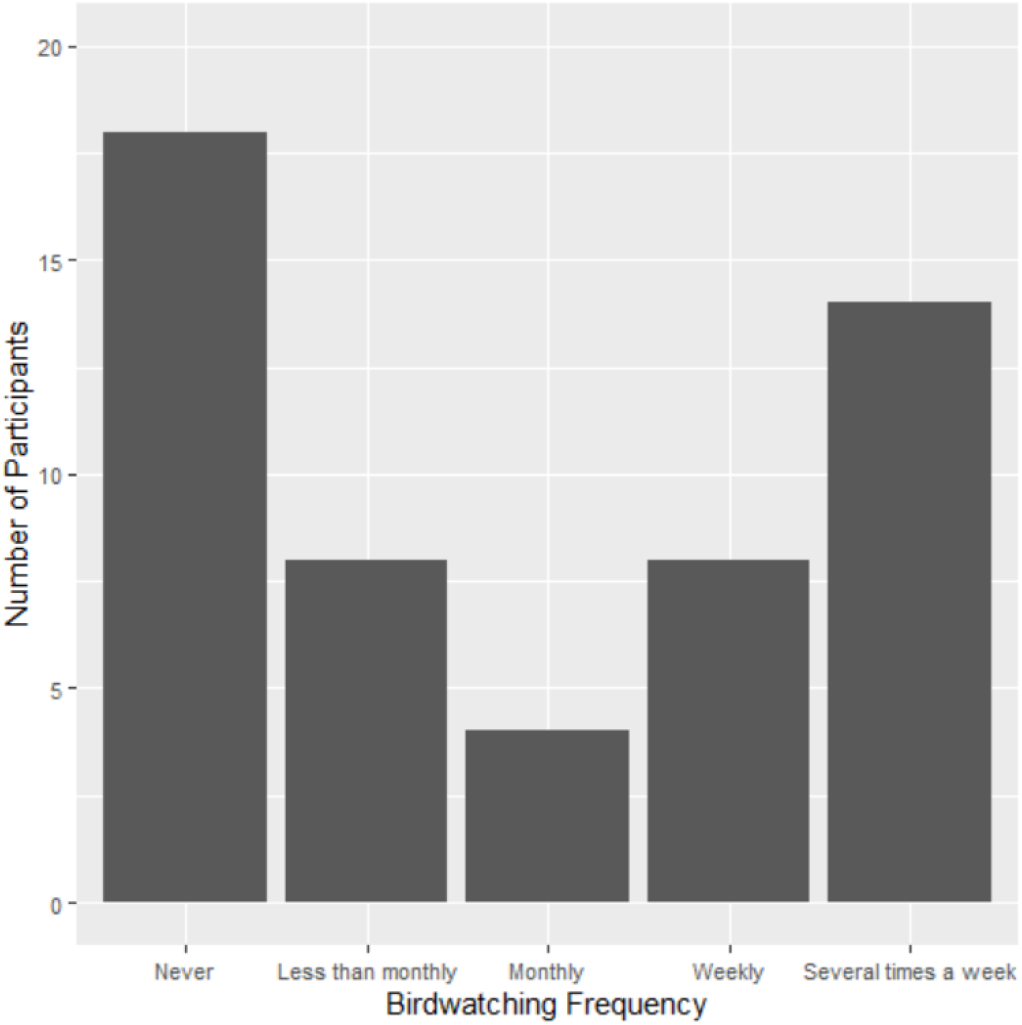
Distribution of birdwatching frequency among participants.

Most volunteers chose the most negative ranking for their opinion about litter (N = 39), the least negative ranking reported being a 6 out of 10. Noticing litter and being around others who litter was also common (see figure 6).

**Figure 6:**
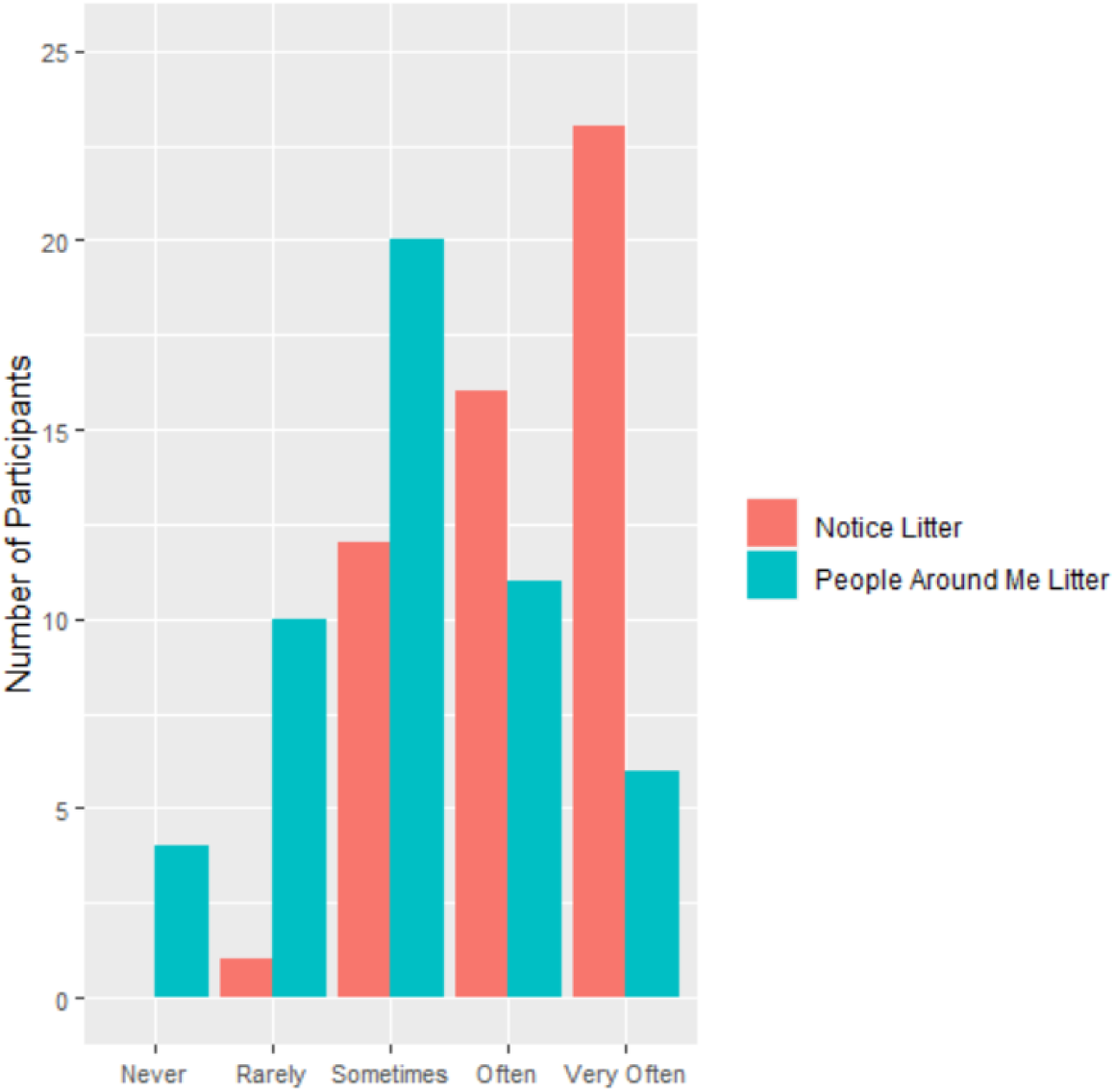
Frequency at which respondents notice litter in their surroundings (red) and frequency at which other people around them litter (such as friends and family).

## 5. Discussion

Given the small sample of observations we received, we cannot draw broad conclusions about the frequency of litter foraging in different UK regions and species. We therefore chose to only report the observations submitted. It should however be noted that the species reported to consume human food in place of naturally occurring food were consistent with previous research (Palacio, 2019): Urban exploiters such as Gulls, Crows and Pigeons were found to be foraging in diverse habitats and predominantly eating human food in place of naturally occurring foods in more than half of the observations submitted for those species. Gulls were observed to exclusively forage and consume human food in place of natural food. Garden birds, Magpies, Starlings, and Blackbirds were reported to consume human food to a much lower degree, and in the case of the Blackbird not at all. These species were also observed foraging in areas such as gardens and parks, as opposed to streets. Although these species relied to a greater degree on naturally occurring foods, in this study we included food items which could also be provided by humans, such as nuts on bird feeders, as natural food.

Citizen science aims to educate and engage individuals from all backgrounds, whilst raising awareness of and solving real world issues, such as climate change and littering (Gura, 2013; Van Noordwijk et al., 2021). In this study, we aimed to engage individuals from a broad range of ethnic and socioeconomic backgrounds in bird watching. We were partially successful in this endeavour: In our sample gender was well balanced, approximately half the respondents were not already engaged in regular birdwatching and almost none of the participants have been involved with the British Trust of Ornithology (BTO) before. 14 participants made use of the option to enter common names for birds in place of species names, which was added to increase accessibility. 10 volunteers were under the age of 30, six of them 18 or younger. However, our goal of reaching respondents from diverse socioeconomic backgrounds was unsuccessful, as only three of the 55 participants were people of colour and the majority of those that participated had a middle-class income and university degree. Having a sample which lacks diversity in citizen science means that individuals from non-white backgrounds miss out on opportunities to develop scientific knowledge and understanding (Brouwer & Hessels, 2019) as we failed to reach them. Improvement to our recruitment methodologies are needed to capture individuals from a variety of background, such that scientific knowledge and skills can be shared with individuals from all ethnic and socioeconomic backgrounds.

We were unable to track individual changes in the perception of birds and litter over time as only four respondents participated multiple times. Repeated participation may be facilitated through questionnaires which capture the necessary information in as few questions as possible. Additionally, data capture forms should be made as convenient as possible, so that participants are more inclined to participate multiple times in the same data collection effort (Pandya, 2012).

While this citizen science study did not achieve a sufficiently large sample to draw broad ecological conclusions, it was successful in reaching and engaging some volunteers new to birdwatching and bird surveys. We hope this report of the successes and failures of our project can be useful in the planning of future citizen science studies.

## 7. Acknowledgements

We are very grateful to all volunteers who have participated in this survey and submitted their observations. We are also thankful for the help we received in distributing this survey, as well as the feedback we have received from Dawn Balmer, Amanda Mead, Kirsty Neller, and Tom Dickins during the early planning stage and the development of the questionnaire.

## 8. Author contributions

S.S. developed the core idea, distributed the survey, analysed the data, and wrote the methods and results sections. W.D.H.W. contributed to the design of the questionnaire, the distribution of the survey, wrote parts of the introduction and discussion. K.C. contributed to the design of the questionnaire, the distribution of the survey and wrote parts of the introduction.

## 9. Appendix A – Questionnaire

1. What bird did you see? You can give the exact name (such as “Carrion crow”), the broad name (such as “crow”), or describe what it looks like (“black bird the size of a football”). If you want to find out which bird you saw you can look for it in the RSPB Bird Identifer.
2. What did the bird eat (e.g. “a worm” or “a cheeseburger”)
3. How long was it eating there?
  a. Less than a minute
  b. One minute
  c. Five minutes
  d. 10 minutes or more
4. Where there any other birds?
  a. Yes, and they looked like my bird (how many other birds?)
  b. Yes, but they looked different than my bird (how many other birds, what did they look like?)
  c. No
5. Did you notice any other behaviour, such as fighting, feeding other birds, singing…?
6. Did the bird finish eating?
  a. Yes
  b. No, it was interrupted (by what?)
7. On what date and at what time did you see the bird?
8. Where in the UK did you see the bird? You can answer with the postcode, location name (e.g. name of the city, park or forest), or with “What3Words”.
9. What type of location was it?
  a. Park
  b. Street
  c. Garden
  d. Forest
  e. Coast
  f. Other (please specify)
10. How old are you?
11. What gender do you most identify with?
12. Have you ever been a BTO volunteer?
  a. Yes
  b. No
13. What is your highest education level?
  a. Secondary
  b. College/Sixth Form
  c. Undergraduate degree (Bachelor)
  d. Graduate degree (Master)
  e. Post-graduate degree (PhD)
  f. None
14. What is your household income?
  a. Less than £20,000 per year
  b. £20,000 to £40,000 per year
  c. £40,000 to £60,000 per year
  d. £60,000 to £80,000 per year
  e. More than £80,000 per year
15. What is your ethnicity?
  a. English/Welsh/Scottish/Northern Irish/British
  b. Irish
  c. Gypsy/Irish Traveller
  d. Other White Background
  e. Indian
  f. Pakistani
  g. Bangladeshi
  h. Chinese
  i. Other Asian Background
  j. African
  k. Caribbean
  l. Other Black/African/Caribbean Background
  m. Arab
  n. Other background not listed here
16. How often do you notice birds in your day-to-day life?
  a. All the time
  b. Most of the time
  c. Sometimes
  d. Rarely
  e. Never
17. How often do you go birdwatching?
  a. Several times per week
  b. Once per week
  c. Once per month
  d. Less than once per month
  e. Never
18. How do you feel about birds?
  a. Like them a lot
  b. Like them a little bit
  c. Like some, dislike others
  d. Neither like nor dislike them
  e. Dislike them a little bit
  f. Dislike them a lot
19. How often do you notice litter in your day-to-day life?
  a. All the time
  b. Most of the time
  c. Sometimes
  d. Rarely
  e. Never
20. How often do people around you litter?
  a. Very often
  b. Often
  c. Sometimes
  d. Rarely
  e. Never
21. How do you feel about litter? (from 0 (doesn’t bother me) to 10 (bothers me a lot))

## 10. Appendix B – Additional Data

**Table 2:** Single observations excluded from grouped analysis.

| **Bird** | **Food Type** | **Food Location** | **Habitat** |
| --- | --- | --- | --- |
| Mallard | Natural | From the ground | Garden |
| Goose | Natural | From the ground | Park |
| Jackdaw | Human | From the ground | Garden |
| Rook | Human | From the ground | Garden |
| Siskin | Natural | Bird feeder | Garden |
| Treecreeper | Natural | Tree trunk | Garden |
| Green Woodpecker | Natural | From the ground | Garden |

**Table 2:**
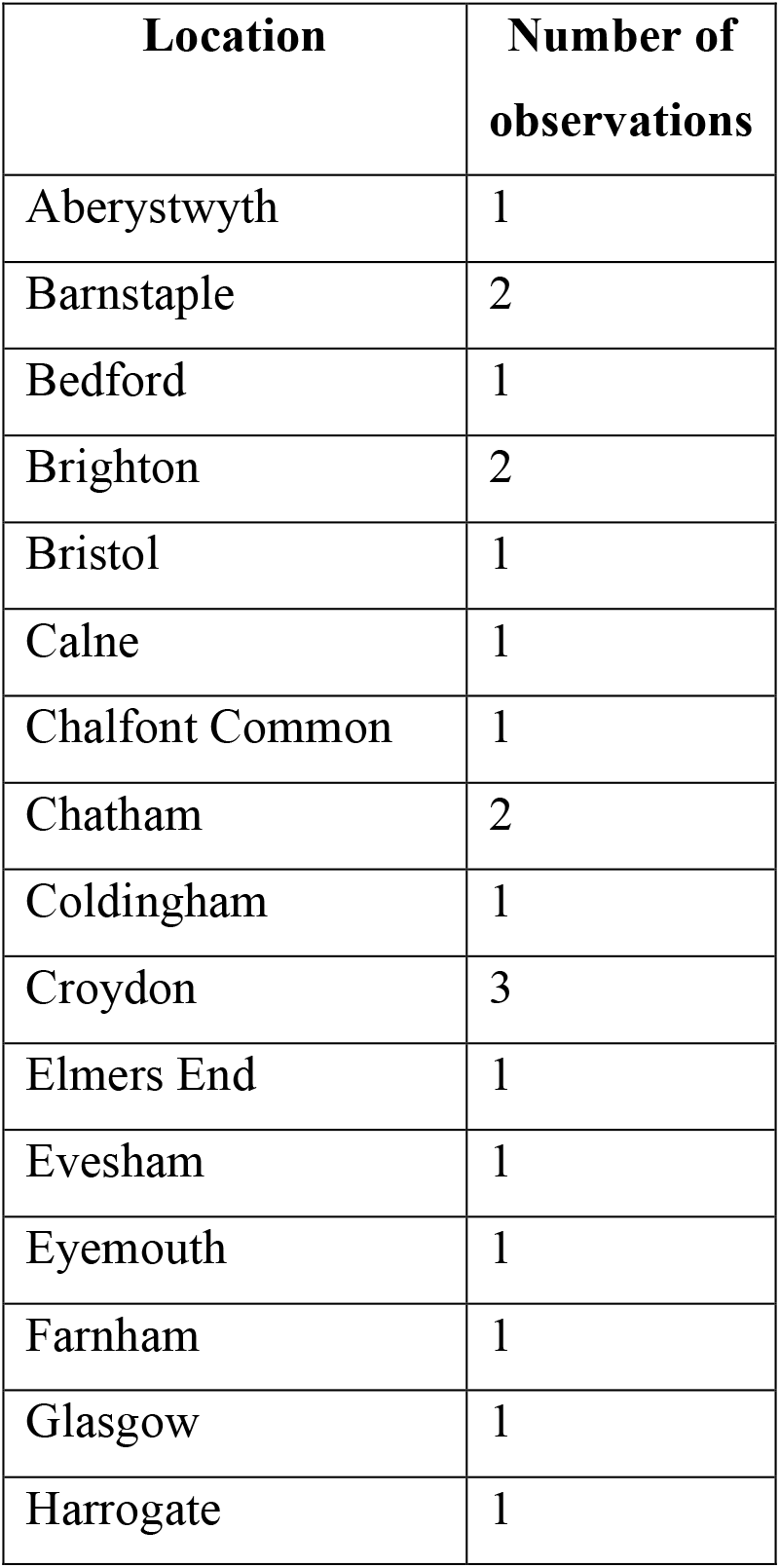
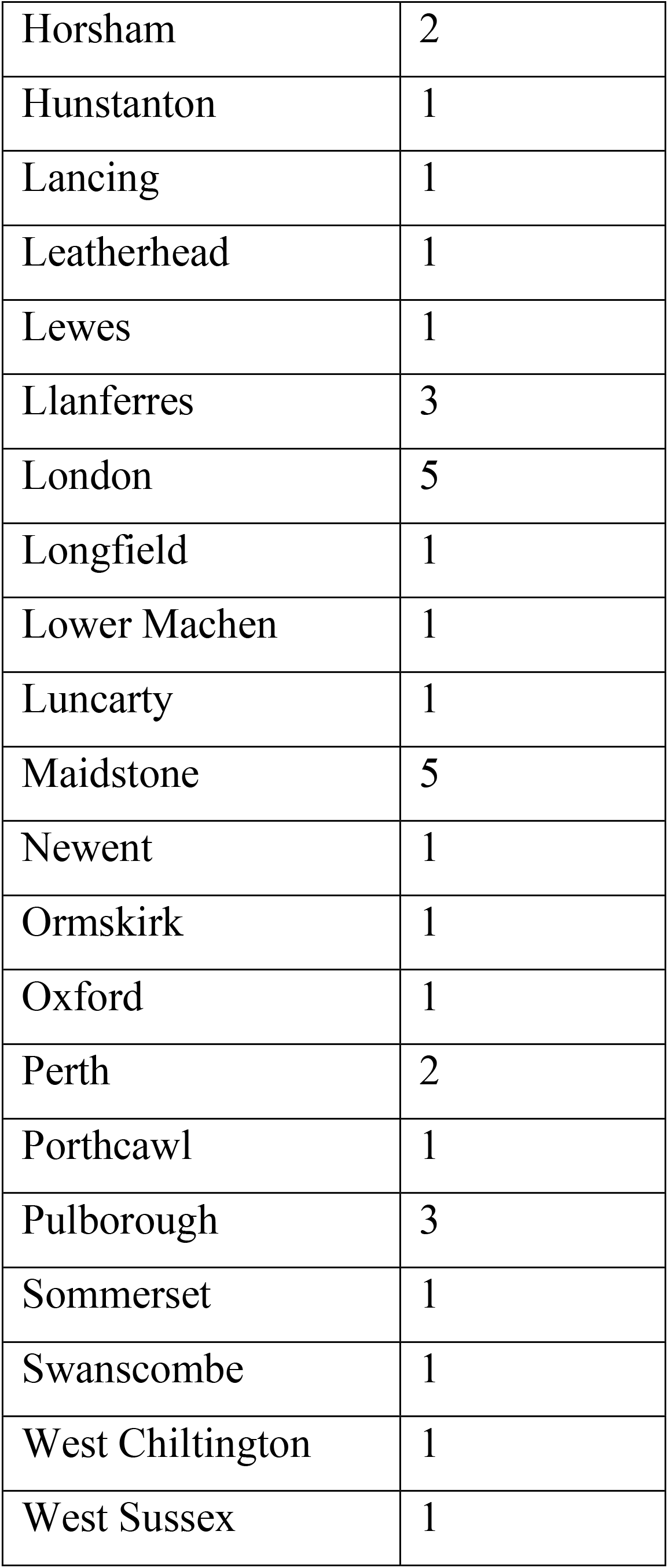
Cities and towns from which observations were reported and included in the analysis.

## Notes

### Competing Interest Statement

The authors have declared no competing interest.

